# High Gamma and Beta Temporal Interference Stimulation in the Human Motor Cortex Improves Motor Functions

**DOI:** 10.1101/2021.03.26.437107

**Authors:** Ru Ma, Xinzhao Xia, Wei Zhang, Zhuo Lu, Qianying Wu, Jiangtian Cui, Hongwen Song, Chuan Fan, Xueli Chen, Junjie Wei, Gongjun Ji, Kai Wang, Xiaoxiao Wang, Bensheng Qiu, Xiaochu Zhang

## Abstract

**Background:** Temporal interference (TI) stimulation is a new technique of noninvasive brain stimulation. Envelope-modulated waveforms with two high-frequency carriers can activate neurons in target brain regions without stimulating the overlying cortex, which has been validated in mouse brains. However, whether TI stimulation can work on the human brain has not been elucidate.

**Objective:** To assess the effectiveness and safety aspect of the envelope-modulated waveform of TI stimulation on human primary motor cortex (M1).

**Methods:** Participants attended three sessions of 30-min TI stimulation at 2 mA during a random reaction time task (RRTT) or a serial reaction time task (SRTT). Motor cortex excitability was measured before and after TI stimulation.

**Results:** In the RRTT experiment, only 70 Hz TI stimulation had a promoting effect on the reaction time (RT) performance and excitability of the motor cortex compared to sham stimulation. Meanwhile, compared with the sham condition, only 20 Hz TI stimulation significantly facilitated motor learning in the SRTT experiment, which was significantly positively correlated with the increase in motor evoked potential.

**Conclusion:** These results indicate that the envelope-modulated waveform of TI stimulation has a significant promoting effect on human motor functions, experimentally suggesting the effectiveness of TI stimulation in humans for the first time and pave the way for further explorations.

## Introduction

The human motor system can quickly react to external stimuli through delicate control of skeletal muscles by the neural activity of the motor cortex [1]. Repetitively performing the same actions in sequential manners allows humans to acquire new motor skills [2]. During these functions, high gamma and beta brain oscillations of the motor cortex play important roles. The functional separation of these two oscillations is an important question to be investigated. Previous studies have found that high gamma brain oscillations are transiently increased during movement and they have a promoting effect on movement initiation[3-5]. Meanwhile, beta activities in the motor cortex are considered an important component of motor learning [6-8]. Modulating these oscillations may be useful to improve motor skills.

Electrical stimulation is the most direct way to regulate electrically oscillating neural activities. Two kinds of electrical stimulation techniques have been extensively used. The first is deep brain stimulation (DBS), which has been proven to be an effective treatment for treating Parkinson’s disease [9, 10]. The delivery of DBS requires invasive surgery, thus presenting the potential for surgical complications [11]. Another method is transcranial electrical stimulation (tES) [12, 13], which can modulate brain activities in noninvasive ways [14-16]. TES applied with alternating current, i.e., transcranial alternating current stimulation (tACS), has been used to facilitate oscillation activity within specific frequency ranges [17, 18]. Many studies have shown that tACS can modulate motor-related oscillation brain activities, which could result in changes in cortical excitability and motor function improvement [19-22]. However, currents of tES applied over the scalp were found to be significantly attenuated when traveling through the skin, subcutaneous soft tissue and skull [23]. Thus, the depth of stimulation is limited. Participants reported side effects during tES stimulation, such as phosphene perception, skin sensations and even skin burns under the stimulation electrode [24-26].

To overcome the limitations of these two electrical brain stimulation techniques, temporal interference (TI) stimulation (Figure 1) has been recently proposed [27], which has caused considerable excitement in the research community [28-33]. This new technique can be applied by delivering two electric fields at frequencies that are too high (≥1 kHz) to elicit neural firing (Figure 1b-c). The frequency difference between these two electric fields is within the range of brain oscillations (e.g., 20 Hz, 70 Hz, etc.), which could result in a prominent envelope modulated electric field (Figure 1d) in a targeted brain region. TI stimulation has been proven to be effective in driving the firing patterns of hippocampal neurons without recruiting neurons in the overlying brain cortex and evoking different motor behaviors in mice [27].

**Figure 1.**
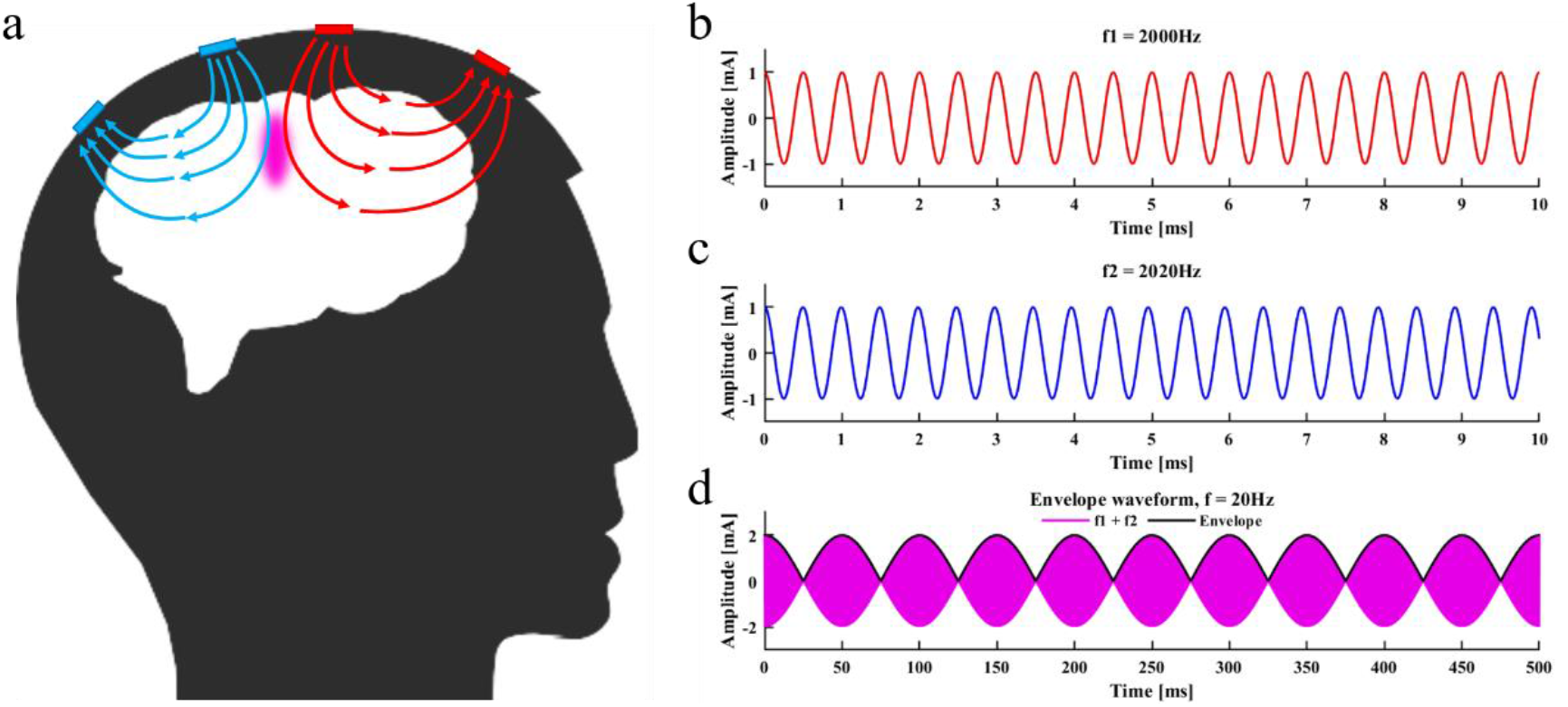
Schematic diagram of TI stimulation. (a) Stimulation of a specific brain region by TI stimulation. (b) One high-frequency (2000 Hz) electric current involved in TI stimulation. c) Another high-frequency (2020 Hz) electric current involved in TI stimulation. (d) Envelope modulated current waveform of TI stimulation, which is generated by the superposition of the two current waves shown in (b) and (c).

Based on the concept of TI stimulation, several modeling and computation studies have been performed to explore the feasibility of TI stimulation in the human brain but have revealed different results. Several studies reported that TI stimulation generated field strength in deep brain regions similar to tACS with smaller electric field intensity in superficial areas [34-38]. A computational modeling study explored the impact of TI stimulation on neuron levels but found that not all types of neurons responded to TI stimulation [39]. Another simulation study reported that TI stimulation might have effects other than activating neurons in the target area, such as conduction block in off-target areas [40].

However, no data about the actual effect of TI stimulation on human brains have been reported thus far. There are many structural differences between the brains of humans and mouse, causing the field generated by TI stimulation in the human brain to not reach the same intensity in the mouse brain [41]. The stimulation waveform of TI stimulation is an envelope-modulated waveform produced by the superposition of two sine waves (Figure 1d), which is much more complex than conventional tES. Whether TI stimulation has a comparable effect with conventional tES on the human brain is unknown. More importantly, whether TI stimulation is safe and tolerable to people is also an urgent issue to explore.

In this study, we implemented TI stimulation targeting the left primary motor cortex (M1) of healthy participants to validate the effectiveness of TI stimulation on the human brain. Considering the prior investigations of oscillations related to M1, we designed two stimulation conditions with envelope frequencies of 20 Hz (beta) and 70 Hz (high gamma). A sham condition was used as a control. To explore the influence of TI stimulation on different levels of motor functions, two motor tasks were employed, including a random reaction time task (RRTT) and a serial reaction time task (SRTT). RRTT is a single reaction time task, and the order of the reactions is totally randomized. SRTT contains repeatedly recurring response sequences, which can be learned by participants [42]. Due to the distinct functions of high gamma and beta oscillations in the human motor cortex, we hypothesized a promotion of reaction speed induced by 70 Hz TI stimulation in RRTT and a more significant effect of 20 Hz TI stimulation on motor learning in SRTT. We also measured motor cortex excitability before and after TI stimulation using an input-output (IO) curve or motor evoked potential (MEP) elicited by single pulse transcranial magnetic stimulation (TMS) [43, 44], which was hypothesized to be facilitated by TI stimulation based on previous findings [19, 21, 45, 46].

## Material and Methods

### Participants

We recruited 27 healthy adult volunteers in the RRTT experiment, and 6 participants were removed from the analysis because of technical issues (a decrease in current due to poor contact and current crosstalk due to the flow of conductive paste). Data from the remaining 21 participants were included in the analysis (6 females, mean age ± SD: 22.429 ± 2.249 years, mean education level ± SD: 15.762 ± 2.166 years, mean handedness score ± SD: 86.667 ± 17.127). Another 33 healthy adults volunteered to participate in the SRTT experiment, but 1 participant was removed due to the sliding of electrodes, 1 participant was rejected because he switched his performing hand, and 2 participants’ data were removed because of technical issues (current crosstalk due to the flow of conductive paste). Therefore, 29 participants remained to be analyzed in the SRTT experiment (15 females, mean age ± SD: 22.103 ± 2.024 years, mean education level ± SD:15.966 ± 1.991 years, mean handedness score ± SD: 77.672 ± 23.792).

All participants reported no history of craniotomy or injury to the head, no personal or family history of neurological or psychiatric disease, no metal implants or implanted electronic devices, no skin sensitivity and no use of medicine during the experiment. For safety reasons, any participant who was pregnant or could be pregnant was rejected. All participants were right-handed as assessed using the Edinburgh handedness inventory [47] and had normal or corrected-to-normal vision. Informed consent was obtained prior to any involvement in the study. This study was approved by the Human Ethics Committee of the University of Science and Technology of China (IRB Number: 2020KY161).

### Experimental design

The experimental procedures were the same for both the RRTT and SRTT experiments except for the detailed motor tasks and MEP procedures. At the beginning of the procedures, individual M1 location was identified by single pulse TMS, and baseline motor cortex excitability was measured. Before stimulation, the participants were asked to perform a practice task with 24 random button presses. Formal experimental tasks (RRTT or SRTT) started 10 minutes after the beginning of TI stimulation. After the 30-minute stimulation, motor cortex excitability was measured again to detect the change in excitability of M1 (Figure 2a). Participants visited the laboratory three times, at least three days apart, to avoid any influence of the carry-over effects of stimulation.

**Figure 2.**
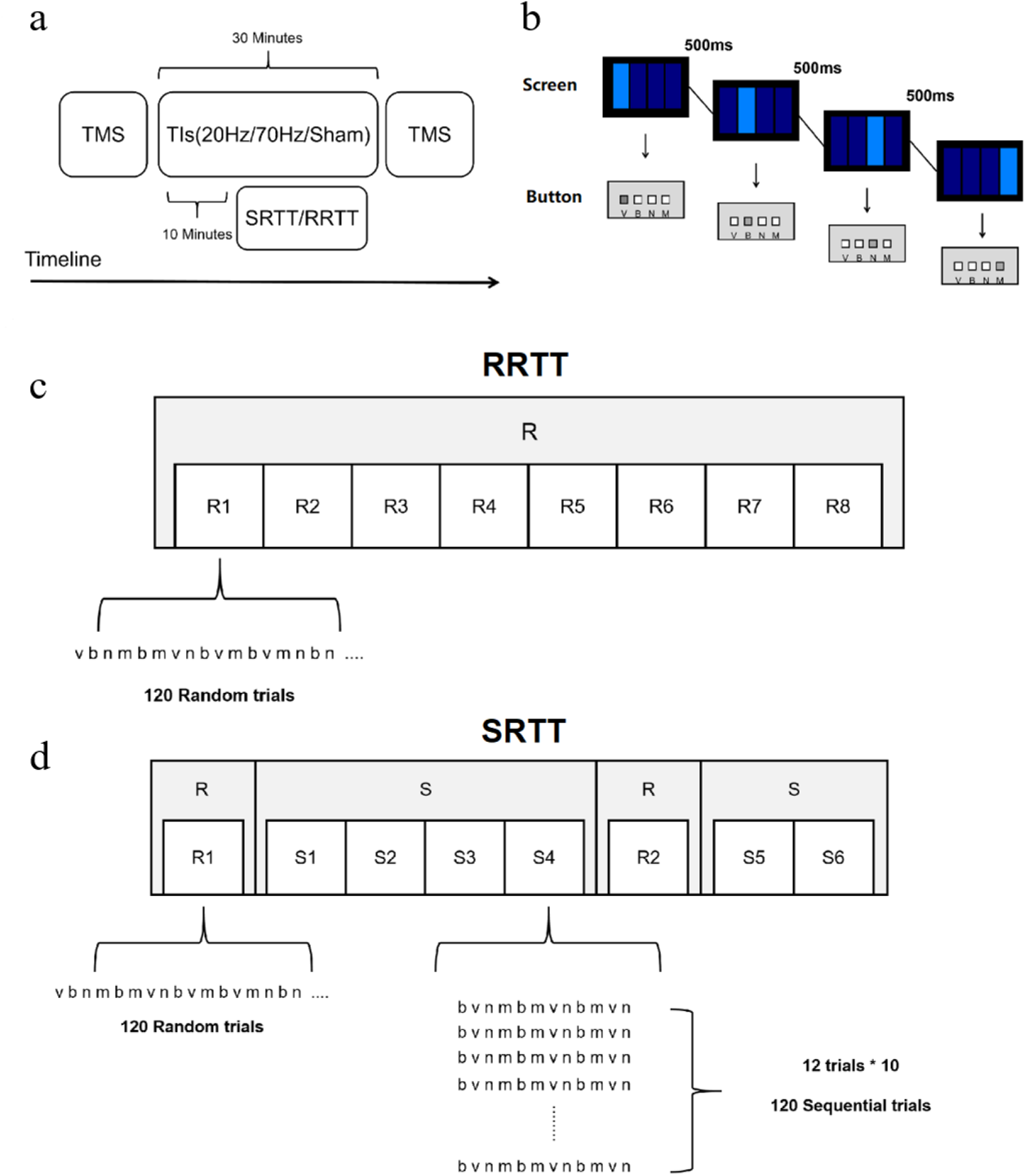
Experimental design and motor tasks. (a) The experimental procedures. (b) Motor task implemented in our experiments. (c) RRTT. (d) SRTT. In R blocks, there were three 12-item (bnmvnbmnvbvm, nvnmbvmnbmvb, mvbmnbnvmbvn) sequences with comparable difficulty for each experimental session in a counterbalanced way.

### TMS and MEP

Single pulse TMS was delivered manually using a 70-mm air-cooled figure-eight coil and a Magstim Rapid^2^ stimulator (The Magstim Company Ltd., Whitland, United Kingdom) with the navigation system of Brainsight (Brainsight, Quebec, Canada). The coil was placed tangentially over the scalp of the left hemisphere to search for the hotspot of the right first dorsal interosseus (FDI), which was marked on a medical elastic bandage on the participants’ heads after the search process. The handle of the coil was pointing posterolaterally 45° from the midline [48, 49]. An electromyogram (EMG) of the right FDI was recorded by a pair of Ag-AgCl electrodes in a belly tendon montage using the EMG module of the navigation system. The resting motor threshold (RMT) was defined as the lowest stimulus intensity that could elicit an MEP in the resting muscle with an amplitude of 50 μV (peak-to-peak) or greater in at least 5 out of 10 recordings [44, 50].

In the RRTT experiment, we applied 15 pulses over the FDI hotspot with an interval of 7 seconds at stimulation intensities of 120%, 100%, 130%, 110% and 140% of RMT before and after TI stimulation [51]. We measured 30 MEPs at a stimulation intensity of 120% of the RMT in the SRTT experiment. Only 120% RMT was used because this intensity corresponds to the linear increase range of the IO curve and is sensitive to the change in M1 excitability [44].

### Motor tasks

The motor tasks were both modified from a SRTT task, which was previously involved in tACS experiments [22, 52]. Participants were instructed to press one of four buttons (V, B, N, M) on the keyboard as fast as possible, according to the position of the light rectangles shown on the screen (Figure 2b). The stimulus remained on the screen until the correct response was made. After 500 ms, a new stimulus was displayed. Eight blocks were included, with 120 trials in each block. The locations corresponding to the light rectangles were pseudorandomly distributed in all 8 blocks (R) in RRTT (Figure 2c). The only difference between SRTT and RRTT was that the reactions were not randomized in some SRTT blocks (Figure 2d). The first block and the sixth block were R blocks. In the remaining blocks, the locations of the light rectangles were repeated in a 12-item sequential manner ten times in each block (S). All of the information about the order of the locations was unknown to the participant, which allowed them to acquire the sequence in an implicit manner. The task presentation and the recording of the reaction times (RT) were conducted using E-Prime 2.0 (Psychology Software Tools, Sharpsburg, USA).

### Temporal interference stimulation

We used five Ag-AgCl electrodes with a radius of 1 cm (Pistim electrode, Neuroelectrics, Barcelona, Spain), four of which were stimulating electrodes and one was the ground electrode located on the mastoid behind the participant’s left ear to avoid current accumulations due to safety considerations. The delivery of the current was provided by a customized battery-driven stimulator with strict safety standards (Figure S1, Supplementary 2.1). The stimulation intensity was peak-to-baseline 1 mA in a single channel. The stimulation electrodes were located 30 mm away from the FDI hotspot along the axis of the Fpz-Oz and T3-T4 in the electroencephalography (EEG) 10-20 system (Figure S2, Supplementary 2.2). Three experimental conditions were included in our design: 20 Hz (2000 Hz & 2020 Hz), 70 Hz (2000 Hz & 2070 Hz) and sham. Stimulation started 10 minutes before the motor task and lasted for 30 minutes. For the sham condition, TI stimulation (20 Hz or 70 Hz) only lasted for approximately 60 seconds (30 s ramp up and 30 s ramp down) at the beginning of this procedure.

### Safety aspects

After the TI stimulation, we asked the participants to complete a subjective questionnaire [24, 25] (Supplementary 2.3), which asked them to rate their sensations including itching, headache, burning, warmth/heat, tingling, metallic/iron taste, fatigue, vertigo, nausea and phosphene during the stimulation and on what extent do they think these feelings were relevant with the stimulation.

### Data analysis

All analyses were performed on MATLAB 2018a (MathWorks, Natick, USA). The mean RT of the correct trials of each block in RRTT or SRTT was calculated. Accuracy was not considered a primary measure because of the ceiling effect (Figure S3). Because the calculation of behavior measures needed to integrate the RT of different blocks, any session containing RT of any block beyond 2SD from all participants’ mean RT was removed. The mean RT of all blocks was considered the behavior measure in the RRTT experiment. Motor learning performance (first implicit learning, FIL, Equation (1); second implicit learning, SIL, Equation (2)) was measured as the RT reduction between S blocks and R blocks in the SRTT experiment.

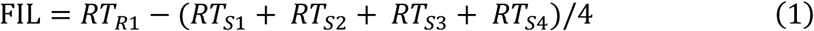

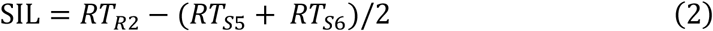

IO curves linearly fitted using the amplitude of MEPs elicited by 100%, 110%, 120%, 130% and 140% RMT were involved in each stimulation condition in the RRTT experiment, and the slope of the IO curve was extracted. Mean MEP amplitudes before and after TI stimulation in each condition were calculated in the SRTT experiment.

Differences in the behavior measures between the stimulation conditions and the control condition (20 Hz vs sham, 70 Hz vs sham) were assessed by two-tailed paired t-tests. Two 2 (condition: 20 Hz vs sham/70 Hz vs sham) × 2 (testing time: before TI stimulation vs after TI stimulation) repeated measures ANOVA was performed on the slopes of the IO curve and MEP amplitudes. We set age, education level and handedness score as covariables to control their potential influence to the motor cortex excitability [53-55]. Since there were significant promoting effects found in the behavior measures, slopes of the IO curve and MEP amplitude before and after TI stimulation were compared by one-tailed paired t-tests with the hypothesis that MEPs would also be facilitated by TI stimulation. Correlations between the behavior measures and increases in the IO slopes or MEP amplitudes in each condition were tested by two-tailed partial correlations, with age, education level and handedness score controlled as covariables. Bonferroni correction was used to correct for multiple comparisons.

## Results

### TI stimulation at 70 Hz promoted the reaction time and M1 excitability

In the RRTT experiment, the stimulation condition of 70 Hz showed the lowest mean RT, which was significantly different from the sham condition (t= -2.953, p_corrected_ = 0.019, Cohen’s d = 0.716) (Figure 3a). There was no significant difference in the comparison between the 20 Hz and sham groups (t = -1.199, p_corrected_ = 0.498).

**Figure 3.**
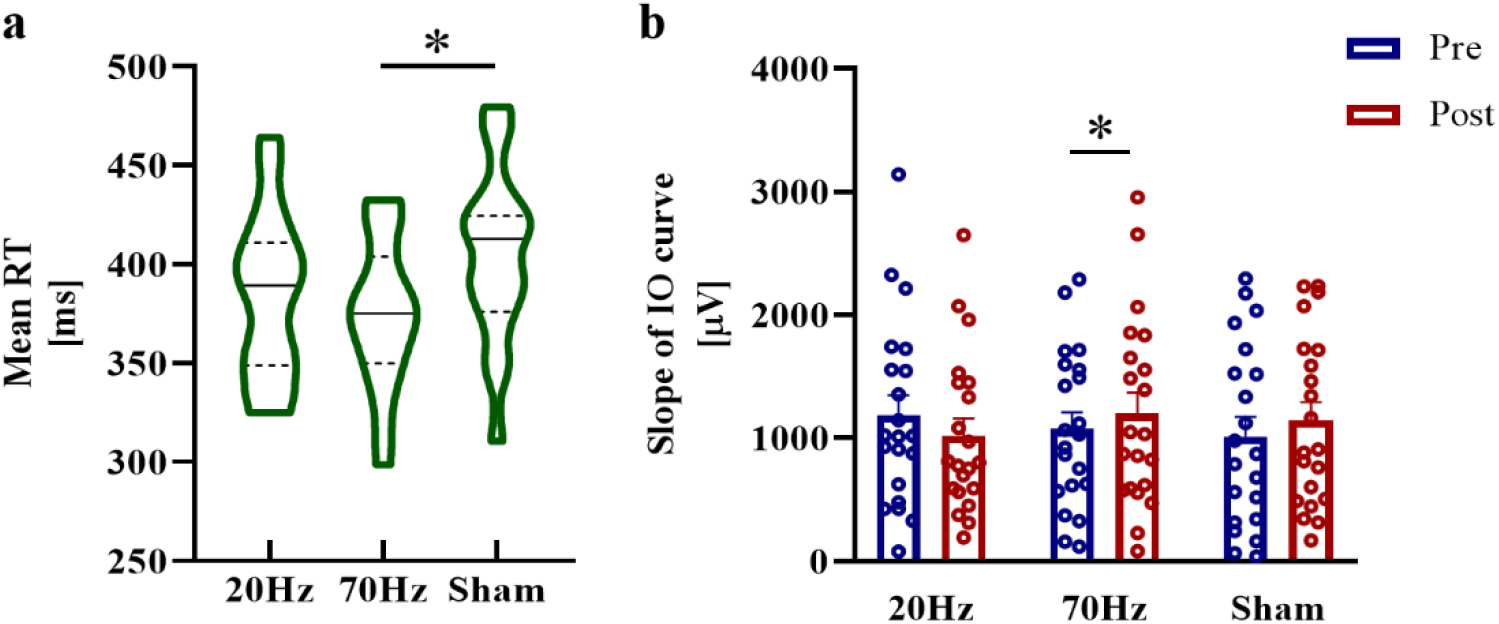
Results of the RRTT experiment. (a) The mean RT of the 70 Hz condition was significantly smaller than that of the sham condition. (b) Significant increases in the IO slope after TI stimulation were found only in the 70 Hz condition. Error bars represent SEM; *significant at p_corrected_ <0.05.

For the slope of the IO curve, we found no significant results either in the main effects of condition or the main effects of testing time or the interaction of the comparison between 70 Hz or 20 Hz with sham (all ps > 0.05). Paired t-tests revealed a significant increase in the IO slope after TI stimulation at 70 Hz (70 Hz: t = 2.395, p_corrected_ = 0.040, Cohen’s d = 0.523, one-tailed) but not at 20 Hz or in the sham condition (20 Hz: t = - 1.075, p_corrected_ = 0.443, one-tailed; sham: t = 1.597, p_corrected_ = 0.189, one-tailed) (Figure 3b).

### TI stimulation at 20 Hz improved implicit motor learning and MEP amplitude

In the SRTT experiment, TI stimulation at 20 Hz showed the highest RT reduction in FIL, which was significantly different from the sham condition, while another comparison did not show significance (20 Hz vs sham: t = 2.577, p_corrected_ = 0.041, Cohen’s d = 0.625; 70 Hz vs sham: t = 0.197, p_corrected_ = 1) (Figure 4a). No significant differences were found in SIL between the stimulation conditions and sham conditions (20 Hz vs sham: t = 0.5116, p_corrected_ = 1; 70 Hz vs sham: t = 1.5716, p_corrected_ = 0.269). For MEP amplitude, when comparing the 20 Hz condition and sham condition, repeated measures analysis of variance (ANOVA) revealed a significant main effect of testing time (F = 4.230, p = 0.050, η^2^ = 0.145), while the main effect of condition (F = 1.463, p = 0.238) and the interaction (F = 0.345, p = 0.563) were not significant. There was also a significant main effect of testing time (F = 6.523, p = 0.017, η^2^ = 0.207) in the comparison between 70 Hz and sham, and no significant result was found in the main effect of condition (F = 2.028, p = 0.167) or in the interaction (F = 0.942, p = 0.341). MEP amplitudes increased after 20 Hz TI stimulation compared with MEP measured before stimulation at a marginally significant level (t = 2.137, p_corrected_ = 0.062, Cohen’s d = 0.397, one-tailed) (Figure 4b). The increase in MEP amplitudes in the 70 Hz and sham conditions was not significant (70 Hz: t = 1.570, p_corrected_ = 0.192; sham: t = 1.254, p_corrected_ = 0.330).

**Figure 4.**
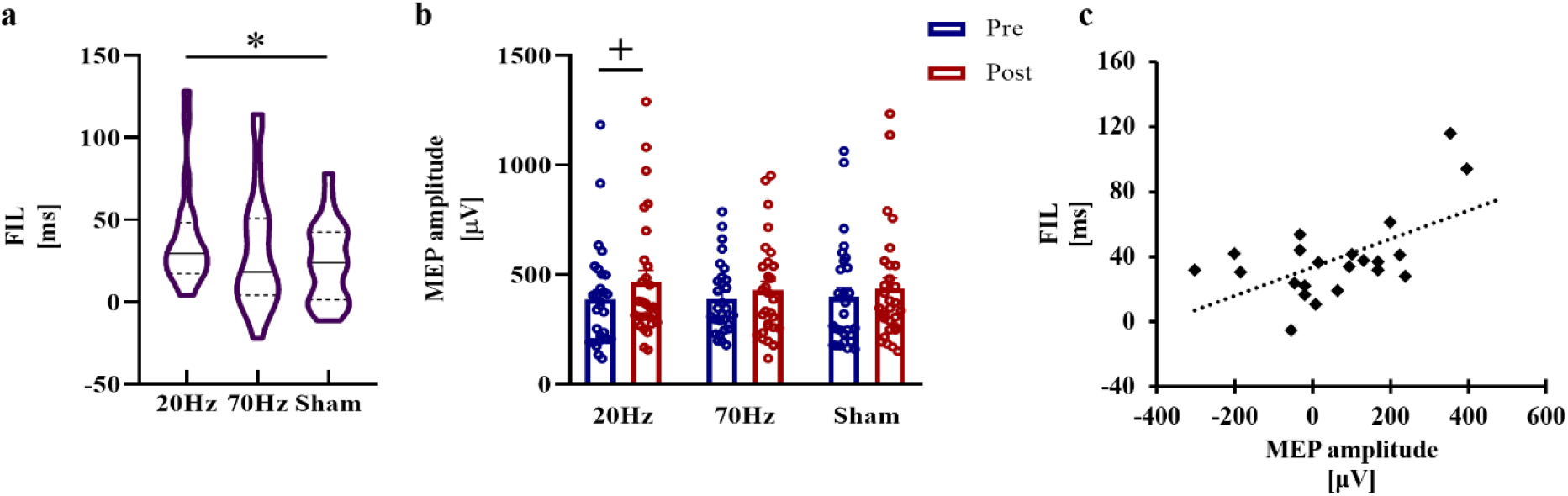
Results of the SRTT experiment. (a) Implicit motor learning during FIL in SRTT. A significant RT reduction was only obtained in the 20 Hz condition. (b) Marginally significant increases in MEP amplitude after TI stimulation at 20 Hz. (c) RT reduction of FIL and the MEP increase in the 20 Hz condition was significantly positively correlated in the 20 Hz condition. Error bars represent SEM; + marginally significant at 0.05< p_corrected_ <0.1, *significant at p_corrected_ <0.05.

The significant reduction in RT during FIL in the 20 Hz condition was positively correlated with the MEP increase (r = 0.580, p_corrected_ = 0.027) (Figure 4c), while RT reductions in the other two conditions showed no significant correlations with the MEP increase (70 Hz: r = 0.073, p_corrected_ = 1; sham: r = -0.360, p_corrected_ = 0.426).

### TI stimulation caused minor side effects on participants

Side effects occurring during TI stimulation were minor and tolerable according to the participants’ descriptions and our observations. Details are given in Table 1 and Table 2. Notably, all discomforts during the sham sessions occurred in the middle of the session or at the end of the session, which could imply that sham stimulation did not directly cause the sensations. Our subsequent investigations of the participants also reported no other side effects after the experiments.

**Table 1.** Discomforts in RRTT

| Sensations | Stimulation sessions | Sham sessions | Percentage |
| --- | --- | --- | --- |
| None | 41 | 19 | 95.238% |
| Fatigue | 0 | 1 | 1.587% |
| Vertigo | 0 | 1 | 1.587% |
| Headache | 1 | 0 | 1.587% |

**Table 2.**
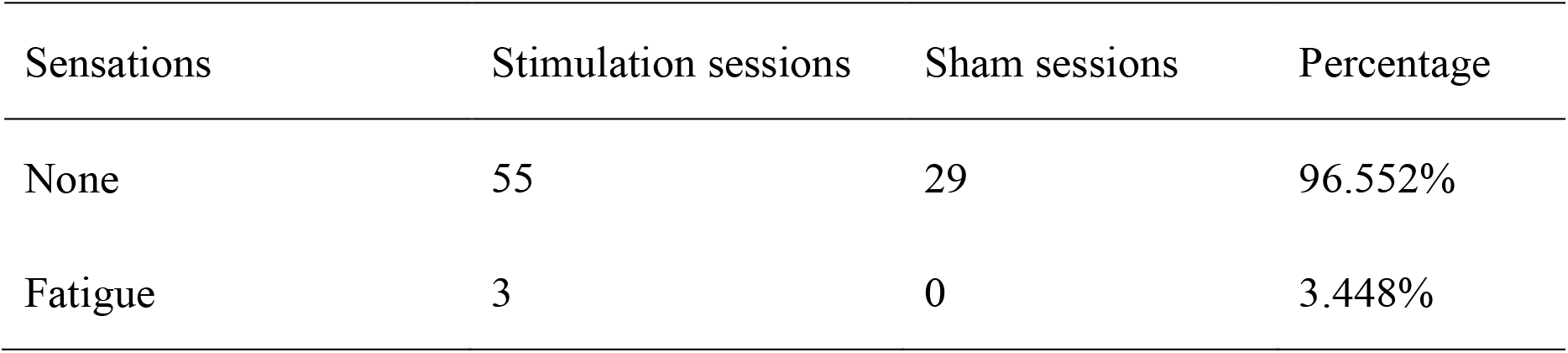
Discomforts in SRTT

## Discussion

In this study, we applied TI stimulation to healthy human participants to explore the modulatory effects of TI stimulation. We investigated the changes in motor performance resulting from TI stimulation applied over M1 in two experiments involving different motor tasks. TI stimulation with an envelope frequency of 70 Hz promoted the RT performance of the motor task compared with the sham condition in the RRTT experiment. TI stimulation with an envelope frequency of 20 Hz applied over M1 enhanced the FIL performance compared with sham stimulation, and the performance was positively correlated with the MEP increase in the SRTT experiment.

### TI stimulation is effective in the human motor cortex

Our study, for the first time, suggests that the idea of TI stimulation is plausible, not only in computational models and experiments on mice[27, 34-38], but also in actual experiments performed on healthy human participants. Since the idea of TI stimulation has been raised, the only in vivo investigations have been performed on mouse brains [27]. The human brain is much larger, and the layers around the brain in humans are thicker, which causes up to 100 times weaker electric fields in the human brain than in the mouse brain at the same stimulation intensity [41]. The stimulation waveform of TI stimulation is an envelope-modulated waveform produced by the superposition of two sine waves, which has not been previously tested on humans. Envelope-tACS using only envelope waveforms of speech without carrier waves have been used to improve speech perception and processing [56-58], but the effects are still controversial [59-61].

Amplitude-modulated tACS (AM-tACS) was proposed as a promising way to allow for effective magnetoencephalography (MEG) or EEG signal reconstruction during electrical stimulation [62-65]. However, similar to TI stimulation, studies on AM-tACS have also mostly focused on simulations, and no systematic experimental test to validate the effectiveness of AM-tACS on humans has been performed. Whether envelope modulated waveforms have comparable effects to conventional tACS is unknown.

To solve these problems, we applied TI stimulation to the human M1 area and found some significant effects. In the RRTT experiment, only 70 Hz TI stimulation promoted RT performance and motor cortical excitability. In the SRTT experiment, only 20 Hz TI stimulation increased the first implicit motor learning and MEP amplitudes. We found significant main effects of testing time in the analysis of MEP amplitude, which might indicate a training effect of the motor learning task [66, 67]. But only the correlation between FIL and MEP increase in 20 Hz condition was significant, indicating the increase of motor cortex excitability related with TI stimulation only occurred in 20 Hz TI stimulation. This can be supported by a meta-analysis, which shows that exogenously applied electric fields in beta frequency range can increase motor cortex excitability [68].

Anyway, to be honest, the effect of TI stimulation shown in this study is not as phenomenal as that in the mice study. Future studies could explore the mechanisms of TI stimulation at the level of brain regions and networks by corresponding neuroimaging techniques, e.g., functional magnetic resonance imaging (fMRI), and try to build a more effective TI stimulation system for human with better understanding to it.

### High gamma and beta oscillations may represent different motor functions in M1

Distinct effects of 20 Hz and 70 Hz TI stimulation may indicate different functions of these two motor cortical oscillations. High gamma and beta are considered vital neural rhythms corresponding to the activation of M1[3-8]. TACS (70 Hz) has been reported to increase motor velocity and motor acceleration during stimulation in visually guided motor tasks [20, 69]. Meanwhile, tACS at 20 Hz has been reported to improve the performance of implicit learning of SRTT in previous studies [22, 52]. These findings could imply that beta and high-gamma neural rhythms predominate in different motor functions in M1. Our results duplicate the functional separation between brain oscillations at 20 Hz and 70 Hz. The functional separation between 70 Hz (2000 Hz & 2070 Hz) and 20 Hz (2000 Hz & 2020 Hz) TI stimulation also supports the hypothesis that electric fields of high-frequency carriers (2000 Hz) may have little contribution to the results because of the intrinsic feature of the neural membrane that filters electrical signals in a low-pass manner [70, 71]. Additional studies could explore the effects of carrier frequency and envelope frequency more deeply.

### The application potential of TI stimulation as a noninvasive brain stimulation technique

We assessed the side effects of TI stimulation by subjective reporting of the participants. In most sessions (>95%), participants reported no side effects. No sensations related to the skin, such as tingling, itching and burning, were reported, and no burns or reddening of the skin were observed by the experimenters. Only fatigue, vertigo and headache were reported in several sessions, including two sham sessions. The side effects reported by participants in this study were far less than those reported for conventional tES [24-26], which indicates that TI stimulation may have advantages over conventional tES in safety, user-friendliness and blinding performance.

Our study indicates that TI stimulation can be used as a new technique to modulate human neural activities in a noninvasive way. We speculate that TI stimulation could be a feasible tool for exploring distinct roles of different brain oscillations in various cognition tasks, especially those neural activities originating from deep brain regions. We preliminarily explored the effects of TI stimulation on human M1. Stimulation effects on other deeper brain regions with more sophisticated functions rely on a better understanding of the working mechanisms and prospects of TI stimulation in humans, which needs to be explored in additional research utilizing combinations of neuron models, finite element modeling simulations and experiments [34]. It has been speculated that regions that are deep but not too small when considered as a fraction of the total tissue volume (e.g., those in stroke, obsessive-compulsive disorder, epilepsy, depression, and spinal cord injury) may be attractive initial indications [33]. Future studies could explore the effect of TI stimulation in deep brain regions and promote the applications of TI stimulation in clinical practice.

## Conclusion

Our study reveals the promoting effect of TI stimulation on human motor functions and motor cortex excitability. TI stimulation with different envelope frequencies showed separate promoting effects on different motor tasks, which implied that TI stimulation may work through a low-frequency envelope. Future investigations of TI stimulation in humans could explore stimulation effects in deeper brain regions under the guidance of modeling works. In summary, TI stimulation could be a promising new technique for noninvasive brain stimulation in humans with clinical application potentials.

## CRediT authorship contribution statement

**Ru Ma:** Project administration, Methodology, Software, Formal analysis, Investigation, Data curation, Writing - original draft, Visualization. **Xinzhao Xia:** Methodology, Hardware testing, Formal analysis, Investigation, Data curation, Writing - original draft, Visualization. **Wei Zhang:** Methodology, Software, Writing - review & editing. **Zhuo Lu:** Methodology, Hardware design and implementation, Hardware testing. **Qianying Wu:** Formal analysis, Investigation, Data curation. **Jiangtian Cui:** Methodology, Validation, Hardware testing. **Hongwen Song:** Methodology, Software, Writing - review & editing. **Chuan Fan:** Writing - review & editing. **Xueli Chen:** Writing - review & editing. **Junjie Wei:** Methodology, Techniques of TMS. **Gongjun Ji:** Methodology, Techniques of TMS. **Kai Wang:** Methodology, Resources, Techniques of TMS. **Xiaoxiao Wang:** Methodology, Hardware, Supervision. **Bensheng Qiu:** Methodology, Hardware, Supervision. **Xiaochu Zhang:** Conceptualization, Funding acquisition, Methodology, Writing - review & editing, Supervision.

## Acknowledgements

We would like to thank Prof. Bettina Pollok for her kindly help with programming of the experimental tasks. And we thank Dr. Wei Lu for his help with circuit testing.

## Appendix A

### Supplementary data

Supplementary data to this article can be found online.

