## Supplementary for "High Gamma and Beta Temporal Interference Stimulation in the Human Motor Cortex Improves Motor Functions"

1. **Supplemental Figures**

**Figure S1. Design and test of the TI stimulation device.**

**Figure S2. Simulation of TI stimulation on a human head model.**

**Figure S3. Accuracy of RRTT and SRTT experiments.**

2. **Supplemental Methods**

**2.1 Design and test of the TI stimulation stimulator**

**2.2 Simulation of TI stimulation on a human head model**

**2.3 Subjective questionnaire for safety aspects**

**Supplemental References**

1. **Supplemental Figures**

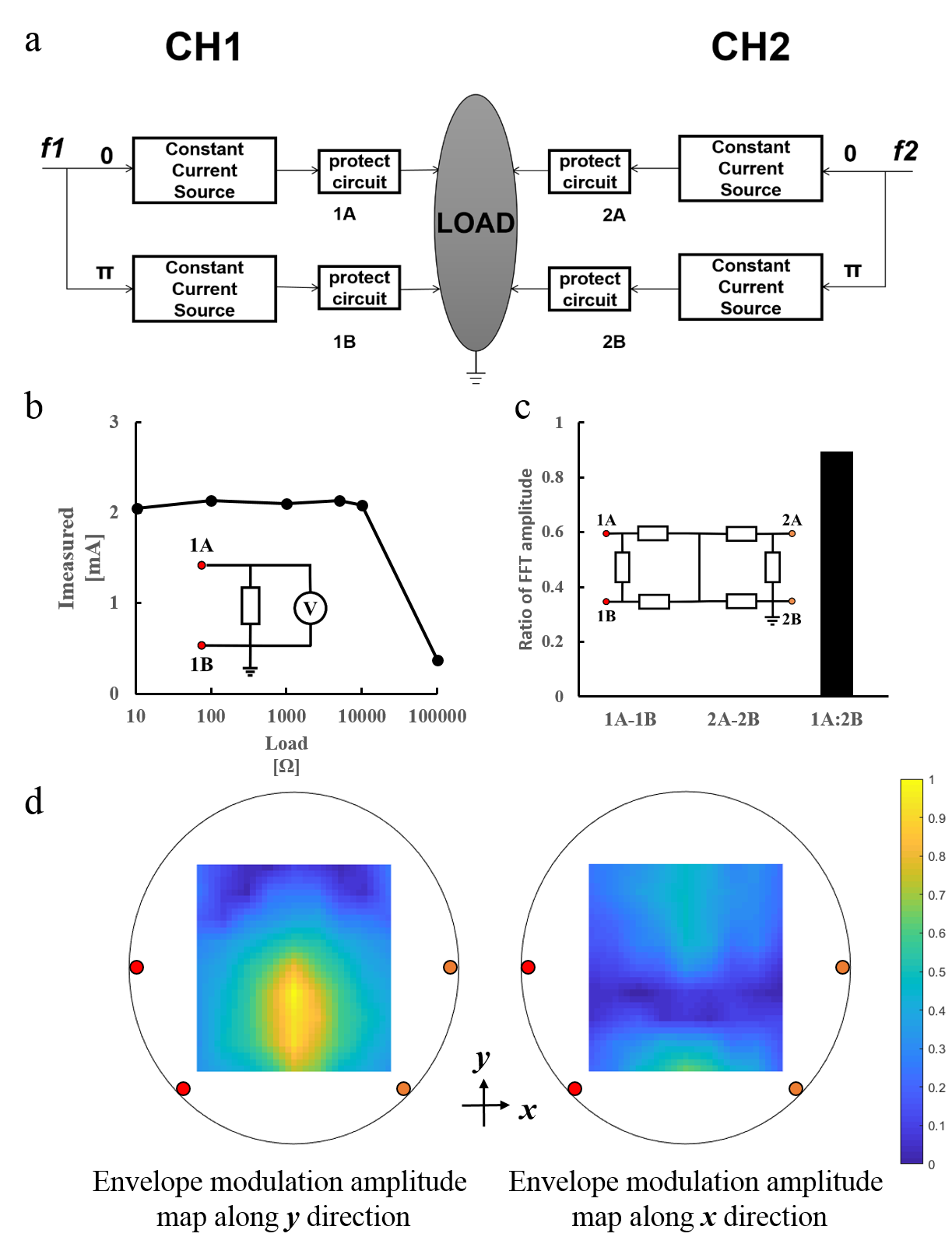

**Figure S1.** Design and test of the TI stimulation device. a) The circuit design of the TI stimulation device. b) The current characterization of the first channel (CH1) of the TI stimulation device on loads with different resistances. c) The characterization of channel isolation of the TI stimulation device. d) The envelope modulation amplitude maps measured on an agar phantom along x and y directions.

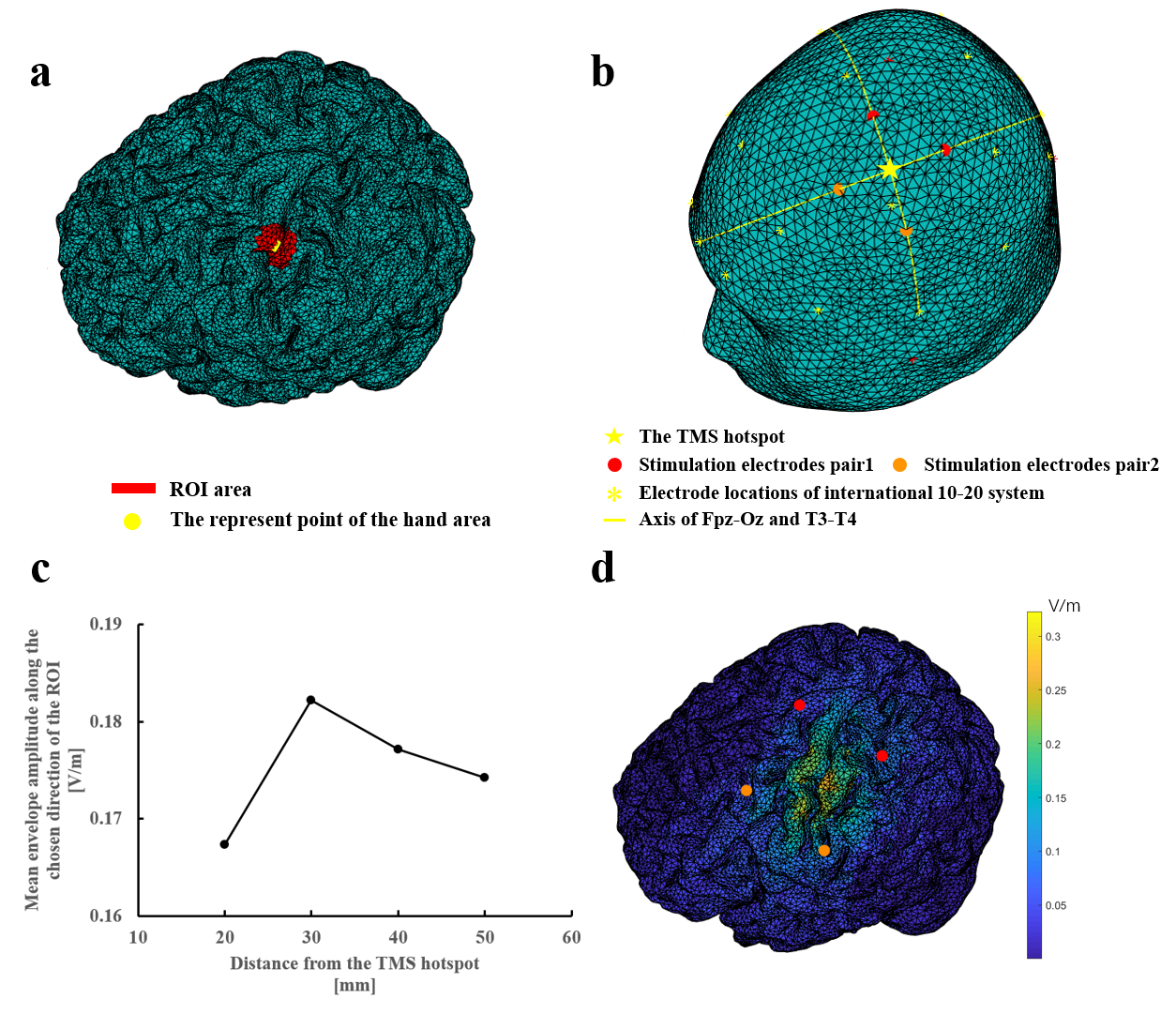

**Figure S2.** Simulation of TI stimulation on a human head model. a) The represent point of the hand area and the region of interest (ROI). b) The stimulation electrodes were located 30 mm away from the TMS hotspot, along the axis of the Fpz-Oz and T3-T4 in the EEG international 10-20 system. c) The mean envelope amplitude of the ROI along the chosen direction was relatively high when the distance from the TMS hotspot and each stimulation electrode was 30 mm. d) The distribution of the envelope amplitude along the chosen direction in the brain with the electrode configuration shown in (b).

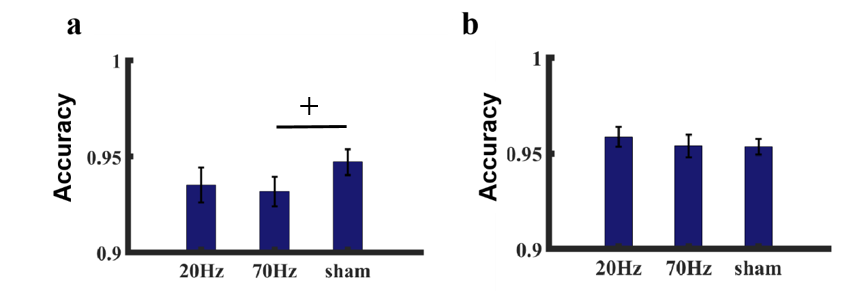

**Figure S3.** Accuracy of the RRTT and SRTT experiments. a) Accuracy of each TI stimulation condition in the RRTT experiment was higher than 93% (mean accuracy ± SD, 20 Hz: 93.513% ± 3.833%; 70 Hz: 93.177% ± 3.267%; sham: 94.705% ± 2.863%). Accuracy in the 70 Hz condition was marginally significantly lower than sham (t = -2.247, p_corrected_ = 0.080), which might reflect the speed-accuracy balance. Differences of accuracy between the 20 Hz condition and sham was not significant (t = -1.915, p_corrected_ = 0.150). b) Accuracy of each TI stimulation condition in RRTT experiment was higher than 95% (mean accuracy ± SD, 20 Hz: 95.850% ± 3.623%; 70 Hz: 95.389% ± 3.671%; sham: 95.335% ± 2.917%). We found no significant differences of accuracy in the SRTT experiment between stimulation conditions and sham. (20 Hz vs sham: t = 0.884, p_corrected_ = 0.774; 70 Hz vs sham: t = 1.080, p_corrected_ = 0.586). Error bars represent SEM; + marginally significant at 0.05< p_corrected_ <0.1.

2. **Supplemental Methods**

**2.1 Design and Test of the TI Stimulator**

TI stimulation device (Figure. S1a) was designed following Grossman et. al.^[1]^ In each channel, two voltage waveforms at frequencies f1 or f2 with a phase difference of π are generated by a DC-powered signal generator (JDS6600, JUNTEX, Zhengzhou, China). The voltage waveforms are then converted into two constant currents outputs. Currents of all the four output ports were monitored by protect circuits to ensure security. Once the amplitude of current at one single output port exceeded the safety threshold (3 mA peak-to-peak), the four output ports would all be cut down by relays. The device is powered by batteries and isolated from mains electricity during use.

We tested the current characterization on different load resistances (Figure. S1b). The output nodes 1A and 1B were connected to loads with resistances between 10 and 100 kΩ. The frequencies of the was 2000 Hz and the current intensity was 2 mA peak-to-peak. Voltage between nodes 1A and 1B was measured using a digital multimeter (UT61E, UNI-T, Dongguan, China) and then converted into current value. Voltages and resistances were measured three times and averaged to reduce noise.

Characterization of channel isolation was tested on a resistance bridge (Figure. S1c). A spectrum analyzer (DSA815, RIGOL, Suzhou, China) was used to measure the frequency spectrum of the currents at the output ports of CH1 between 1A and 1B, the output ports of CH2 between 2A and 2B, and across the resistor bridge between 1A and 2B. The frequencies of the two channels were set as 10000 Hz (f1) and 105000 Hz (f2) and the current intensity of each channel was 2 mA peak-to-peak. Ratio of the FFT amplitude of f1 and f2 was calculated of the three spectrums (Figure. S1c).

We built a phantom to test the electric field distribution generated by the TI stimulation device. The phantom was compounded by 16 g agar powder, 0.2 g sodium chloride and 400 ml water, then heated to boil and poured into a 14 cm diameter petri dish to cool and solidify. A 9 × 9 square-grid was drawn symmetric around the center of the circle on the surface of the phantom. The spacing between grid lines was 1 cm. The frequencies of the two channels were 2000 Hz and 2020 Hz and the intensities of currents were 2 mA peak-to-peak. The envelope modulation amplitude along the ***x*** and ***y*** directions was measured by a digital oscilloscope (DSOX1204G, Keysight Technologies Inc., Santa Rose, USA). The oscilloscope probe terminals were connected to two iron needles located around one point of the grid along ***x*** or ***y*** direction. The distance between the two needles was kept 2 cm. The envelope amplitude was measured on the difference waveform of the two oscilloscope channels.

**2.2 Simulation of TI stimulation on a human head model**

The simulation of TI stimulation was performed using finite element method (FEM) in COMSOL Multiphysics 5.3a (COMSOLAB, Stockholm, Sweden). The electrode montage of TI stimulation was manually optimized to achieved biggest E-field strength along the chosen direction in the target location. The head model was from COMETS toolbox ^[2]^, which was extracted from the standard Montreal Neurological Institute (MNI) brain atlas. The head model was divided into four components and different conductivity values were assigned (scalp: σ = 0.465S/m; skull: σ = 0.01S/m; cerebrospinal fluid: σ = 1.65S/m; brain: σ = 0.276S/m). The chosen direction is pointing posterolaterally at a 45° from the mid-line, which has been widely used in single pulse TMS targeted hand area of motor cortex to approximately make current flow perpendicular to the central sulcus ^[3-5]^,The represent point of the hand area was chosen according to Yousry et al. ^[6]^,and the ROI was defined as a 10 mm radius sphere around it. The TMS hotspot was the projection of this point to the scalp. Electrodes were located equidistant from TMS hotspot along the axis of the Fpz-Oz and T3-T4 of the EEG international 10-20 system.

**2.3 Subjective questionnaire for safety aspects**

The questionnaire [7, 8] asked the participants to rate their discomfort sensations, including itching, headache, burning, warmth/heat, tingling, metallic/iron taste, fatigue, vertigo, nausea and phosphene during the stimulation and on what extent do they think these feelings were relevant with the stimulation. Participants were asked to rate the extent of these sensations that they experienced, from 0 to 4, representing none, mild, moderate, considerable and strong, respectively. Similarly, the relevance to the stimulation was also rated from 0 to 4, representing none, remote, possible, probable and definite, respectively. The influence of those sensations, if any, on the motor task was also evaluated by the participants. The following criteria were used: 1) Only sensations with a score larger than 1 were taken into consideration, and 2) Any sensations that had no relevance at all to the stimulation based on the participants’ descriptions were not considered discomforts elicited by the stimulations.
